# A Novel Method for Testing Antifungal Interactions: Assessing Synergy and Antagonism in *Candida albicans* Clinical Isolates

**DOI:** 10.1101/2024.12.12.628115

**Authors:** Hanna Lundvik, Ramona Santini, Tugce Unalan Altintop, Volkan Özenci, Dan I. Andersson, Nikos Fatsis-Kavalopoulos

**Affiliations:** Department of Medical Biochemistry and Microbiology, Uppsala University; Department of Clinical Microbiology, Karolinska Institutet; Department of Medical Microbiology, Hacettepe University

## Abstract

The rise in antifungal resistance has limited treatment options for serious fungal infections, emphasizing the need for effective combination therapies. However, low-cost and rapid systems to evaluate synergy and antagonism in antifungal combinations are lacking. Here, we introduce a novel in vitro testing method for assessing antifungal interactions in C. *albicans*, enabling the simultaneous testing of three antifungal agents in a single agar plate with overnight results. This method, validated against the checkerboard assay, provides consistent fractional inhibitory concentration (FICi) measurements with reduced variability and workload. We applied this method in a comprehensive screen of 92 clinical C. *albicans* isolates for three antifungals—amphotericin B, fluconazole, and anidulafungin—yielding assessments of a total of 276 distinct combinations of antifungals and isolates. Results revealed isolate-specific interaction patterns, with amphotericin B and fluconazole showing synergy in 1% of isolates, anidulafungin and fluconazole in 19.5%, and amphotericin B and anidulafungin in 23.9%. These findings underscore the need for isolate-specific testing in clinical settings, which this scalable, high-throughput assay can facilitate effectively.

## Introduction

Fungal infections pose a significant health risk, causing life-threatening diseases such as meningitis, pneumonia, and fungaemia, as well as chronic and recurrent conditions like oral and vaginal candidiasis, and asthma (1). The growing threat from fungal pathogens is exacerbated by the increasing prevalence of antifungal resistance, which limits both treatment options and efficacy (2). During the last decades, several resistant fungal species have emerged, such as C. *glabrata, C. parapsilosis*, and *C. auris*, that have very few therapeutic options due to the limited available antifungal arsenal (3–6). Currently, only three major classes of antifungals are routinely used to treat invasive fungal infections, namely; echinocandins, azoles, and polyenes, of the latter only one drug, amphotericin B, is approved for systemic use.

With a limited array of antifungals and the continued rise of resistance, clinicians are increasingly turning to combination therapies to enhance treatment outcomes (7). Combination therapy, used either consecutively or sequentially (3,8,9) is particularly important in treating immunocompromised patients with underlying comorbidities, where monotherapy may not be sufficient for eradication (10).

One antifungal combination, amphotericin B and 5-flucytosine, is already regularly recommended for the treatment of candidiasis (11) and several other combinations are also frequently employed to treat infections (3,8,9). Combination therapy however is not a simple addition of antimicrobial effects. Administering multiple antifungal drugs can lead to synergistic interactions, where their combined action is greater than the sum of their individual effects, or conversely, antagonistic interactions may occur, reducing the overall effectiveness of treatment (11).

In cases of synergistic interactions, the required doses of each drug could be lowered, potentially reducing toxic side effects (12,13). This is especially beneficial given that several antifungal agents are associated with toxicity or contraindications in high-risk patient populations (14). Moreover, the emergence of antifungal resistance could be slowed, preserving the efficacy of available drugs (15) particularly with newly introduced antifungals like ibrexafungerp, where prolonging their clinical lifespan is essential (16).

The potential benefits of identifying synergistic combinations and avoiding antagonistic ones have led to numerous studies aimed at evaluating antifungal interactions (7,17–19). However, the field lacks consensus, with some combinations reported to show either synergistic or antagonistic effects depending on the isolate suggesting that the efficacy of antifungal combinations may be isolate specific (19,20).

In recent years, several high-throughput methods have been developed to predict the combinatorial outcome of antifungal combinations, mainly by utilizing chemogenomic screenings (21–23). However, in vitro methods for studying isolate-specific antifungal interactions have limitations. The checkerboard assay is the most widely used, but it lacks standardization, resulting in significant interlaboratory variability in both performance and data collection (24). Moreover, it is labour-intensive and requires skilled personnel, which may limit its application in resource-constrained settings. Other methods, such as disc diffusion and time-kill assays, are also used but face similar challenges with standardization (25). In contrast, recent advances in antibiotic combination studies have introduced innovative approaches that could be adapted for antifungal combinations (26–29).

In this study, we aimed to develop a reliable and user-friendly method for studying antifungal interactions, with the goal of creating a straightforward pipeline that could lead to much-needed standardization in the field.

Our method builds on an in vitro assay previously used for testing antibiotic interactions in bacteria (26). The CombiANT interaction method, uses a custom culture plate designed to create defined diffusion gradients of three antimicrobials simultaneously. Each antimicrobial is loaded into separate reservoirs surrounding a triangular interaction area. After overnight incubation, distinct zones of growth inhibition appear, allowing for the rapid quantification of drug interactions. Automated image analysis calculates Fractional Inhibitory Concentration indices (FICis) with high accuracy, enabling precise identification of synergistic, additive, or antagonistic effects between antimicrobial pairs.

Here, we optimized and adapted this method into a novel combination plate for testing antifungal interactions, using C. *albicans* as a model organism. This plate allows for the simultaneous testing of three antifungal agents in one agar plate, effectively replacing the need for three individual checkerboard assays. We validated the performance of the combination plate against the commonly used checkerboard assay and applied it to screen interclass antifungal combinations (amphotericin B, anidulafungin, and fluconazole) across 92 clinical C. *albicans* isolates. Through this method validation and its application in the largest screening of clinical C. *albicans* isolates to date, we aim to provide a valuable tool for antifungal combination testing, generate large-scale data on Candida interactions, and facilitate the development of patient-specific combination therapies.

## Results

The primary aim of this study was to develop a method for rapid and straightforward identification of interactions between antifungal agents. To achieve this, we adapted a previously established technique used for detecting interactions between antibiotics targeting bacteria (26,30,31).

Our pipeline is based on a combination plate assay that allows testing interactions between three antifungals simultaneously (Fig. 1). The assay uses a standard petri dish format, incorporating three antifungal-loaded reservoirs (A, B, and C) surrounding a triangular interaction area (Fig. 1I). To prepare the assay, agar infused with antifungal agents is added to the reservoirs and left to solidify. At this stage, the plate remains inert and can be stored refrigerated, limited only by the shelf life of the antifungals. This design allows for the preparation of large batches of plates in advance, which can be used when needed or in large-scale experiments, ensuring consistent handling of reagents.

**Figure 1.**
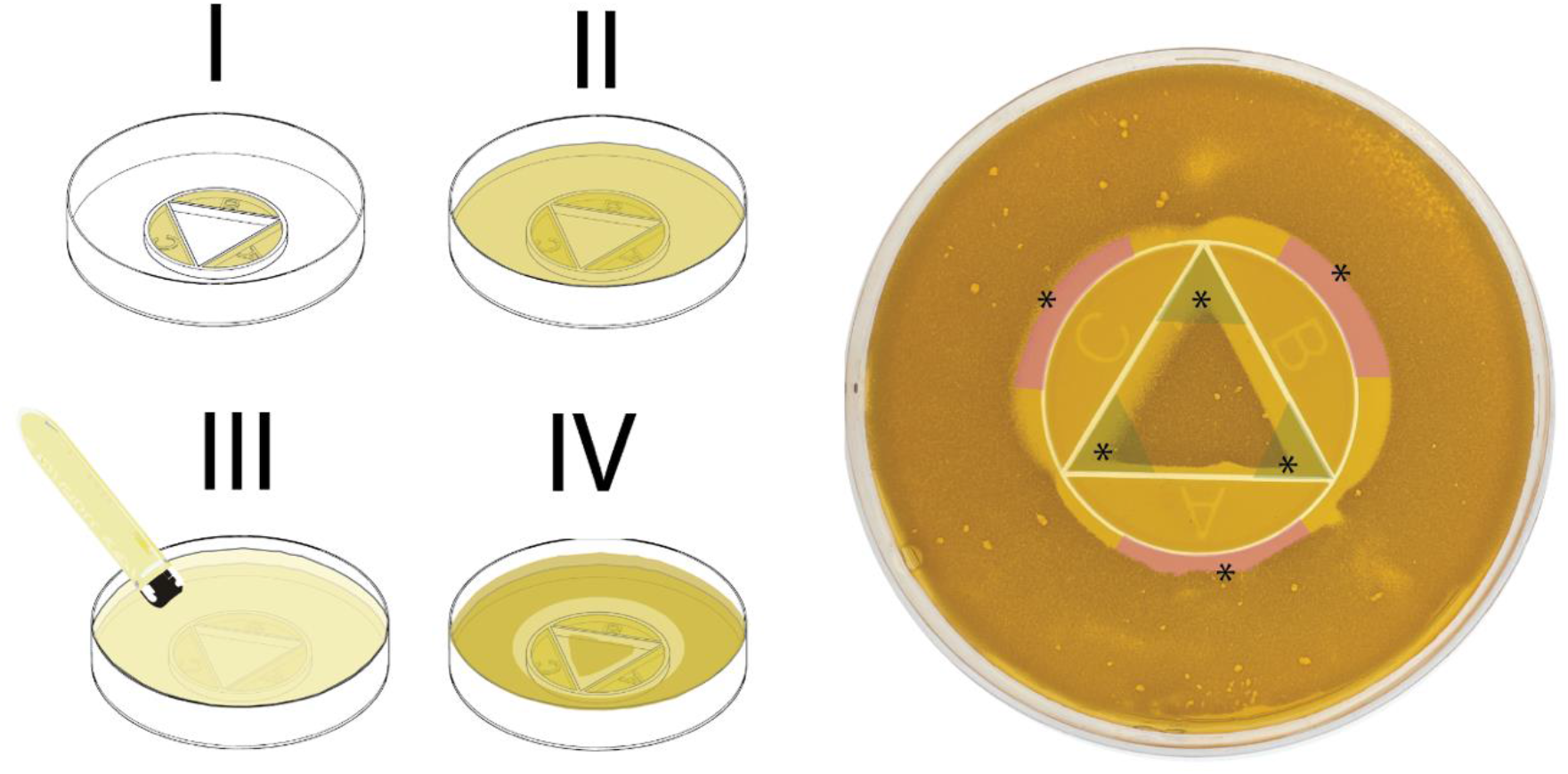
Schematic and workflow of the antifungal combination plate assay for interaction testing. The combination plate assay, designed for testing interactions between three antifungal agents, uses a standard petri dish format with antifungal-loaded reservoirs (A, B, and C) surrounding a triangular interaction area (I). Agar infused with antifungal agents is added to the reservoirs, solidified, and refrigerated until use. The assay is activated by adding a 25 mL agar underlay to allow antifungal diffusion (II), followed by inoculating the plate with a low-temperature gelling agarose overlay containing a fungal inoculum (III). After overnight incubation at 30°C, inhibition zones and growth patterns indicating antifungal interactions are visible (IV). Areas of inhibition for individual antifungals appear (marked in pink), while growth inhibition zones due to antifungal interactions are marked in green. Six key points of interest, marked by asterisks, represent the inhibition edges for each antifungal and the internal growth zone edges near each corner of the interaction triangle, used for calculating Fractional Inhibitory Concentration index (FICi) values for each antifungal pair.

The assay is activated by adding a 25 mL layer of agar to the combination plate, enabling the antifungals to diffuse from the reservoirs (Fig. 1II). After that layer solidifies, fungal isolates are overlaid by inoculating low-temperature gelling agarose, which is spread across the previously solidified agar layer (Fig. 1III). This step is critical for achieving results after an overnight incubation, as the inoculum must contain enough cells to produce readable outcomes despite the longer generation times of fungi compared to bacteria.

Following overnight incubation at 30°C, antifungal interaction patterns specific to each isolate are visible on the plates, with zones of inhibition and growth clearly distinguishable (Fig. 1IV).

On the combination plate, specific points of interest are annotated for analysis. These points include the edges of inhibition zones where individual antifungals act alone (termed inhibitory concentration points, or ICs indicated with asterisks in the pink marked regions of Fig.1 right panel) and the corners of growth zones where combinations of two antifungals produce their combined effects (termed combination inhibitory points, or CPs indicated with asterisks in the green zones of Fig.1 right panel). The ICs are located along the outer boundary of the interaction area, opposite the corresponding antifungal reservoir, marking the point where the antifungal concentration alone is sufficient to inhibit growth. The CPs are located within the interaction area, at the corners of triangular growth zones, representing the combined inhibitory effect of two antifungals.

Using the previously described analysis algorithm(26), these annotated points are aligned with a predefined diffusion model that maps the concentration gradients of the antifungals across the agar. The algorithm calculates the concentrations of the antifungals at both the IC and CP points. Fractional inhibitory concentration index (FICi) values for the three pairwise combinations of antifungals are then calculated using the following formulae and the extrapolated concentrations of the CP and IC points (32):

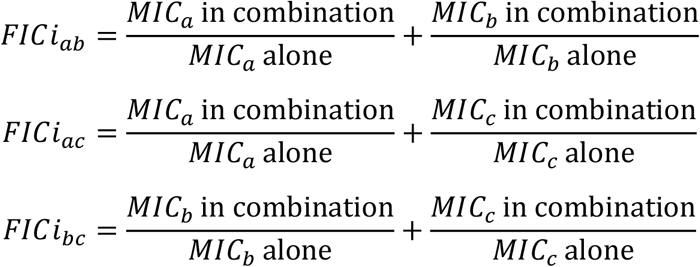

After establishing the protocol, we conducted a large-scale screen for antifungal interactions using a collection of 92 clinical C. *albicans* isolates (Fig. 2). The screening provided FICi values for combinations of fluconazole (FLC), anidulafungin (ANI), and amphotericin B (AMB)— antifungals chosen as representatives of the three main classes used to treat systemic fungal infections. A FICi value of 1 was used as the threshold between synergistic and antagonistic interactions (33,34). Isolates with a mean FICi below 1 were categorized as having synergistic interactions, while those with a mean FICi above 1 were classified as antagonistic (35–38). Our results showed that the AMB-FLC combination exhibited a synergistic interaction in 1% (1/92) of the strains (Fig. 2). The ANI-FLC combination showed synergy in 19.5% (18/92) of the strains (Fig. 2), and the AMB-ANI combination had a synergistic interaction in 23.9% (22/92) of the strains (Fig. 2).

**Figure 2.**
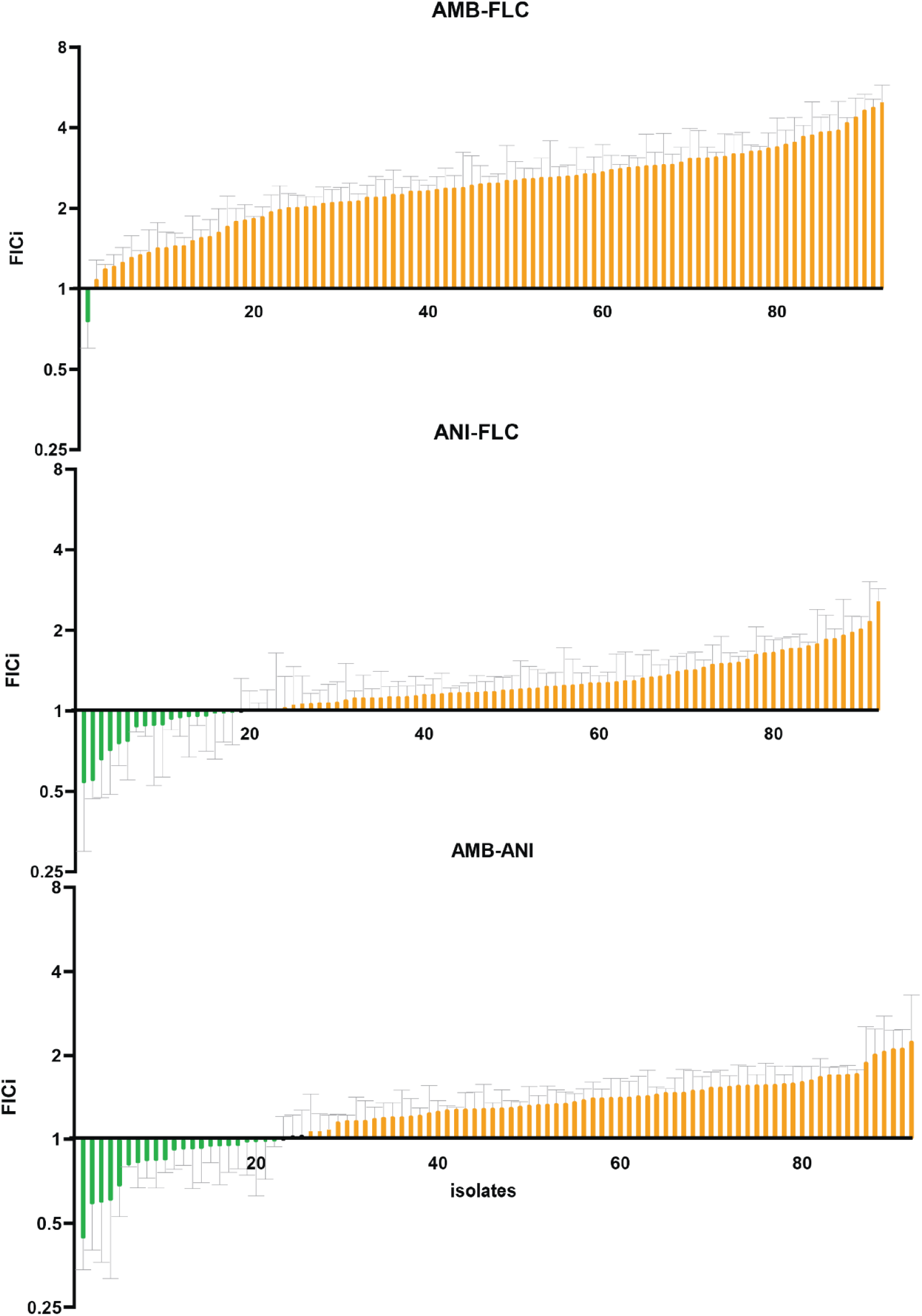
Screening results of antifungal interactions across clinical *Candida albicans* isolates. Results from a large-scale screen of 92 *C. albicans* isolates tested with the combination plate assay for three antifungal pairs,: amphotericin B (AMB) with fluconazole (FLC), anidulafungin (ANI) with FLC and AMB with ANI, shown in ascending FICi order for every combination . FICi values were used to categorize interactions as synergistic (mean FICi < 1,marked in green) or antagonistic (mean FICi > 1, marked in orange). Grey bars mark SEM for n=3.

To verify the accuracy of our method, we compared its results with those obtained using checkerboard interaction assays for the same three combinations (AMB-FLC, ANI-FLC, and AMB-ANI). Checkerboards of AMB-FLC, ANI-FLC, and AMB-ANI were performed for ATCC 90028 C. *albicans* reference strain. FICi values were calculated (32) at 95% growth inhibition for the combinations of AMB-ANI and AMB-FLC and 75% for of ANI-FLC in concordance with the Broth micro dilution (BMD) thresholds for determining the inhibitory effect of each antifungal (39) (supplementary protocol 1). FICi values from the combination plate and checkerboard assays were similar (Fig. 3 left), with a paired nonparametric t-test showing no significant differences between the methods.

**Figure 3.**
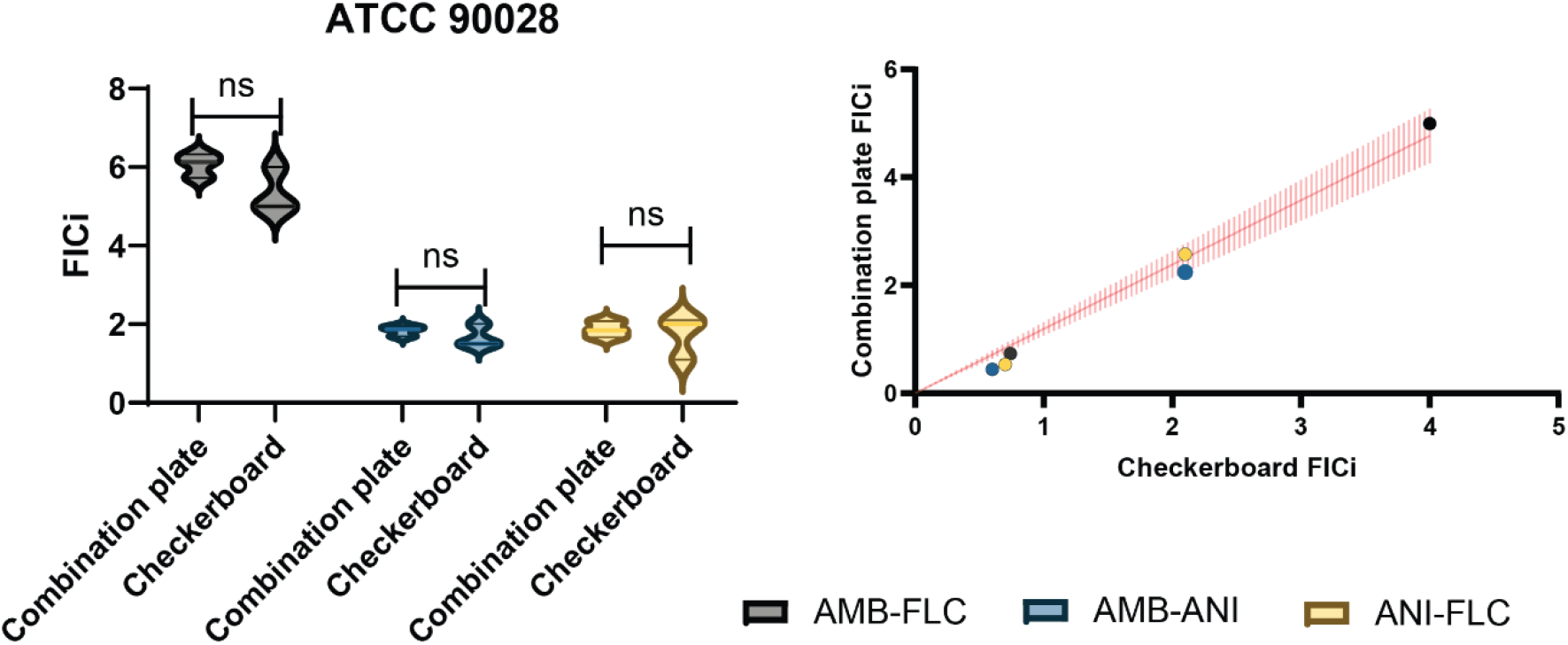
Validation of combination plate results against checkerboard assay for antifungal interaction testing. Left: FICi values of the ATCC 90028 *C. albicans* reference strain with the Combination plate and Checkerboard assay, non-parametric t test shows no significant difference between the two methods. Right: Regression analysis of FICi values obtained from both methods for selected clinical isolates (those with the most synergistic and antagonistic FICi values), regression slope shown in read between 1.066 and 1.317 (95% CI, p = 0.001).

To test the discrepancy between the two methods when applied on clinical strains, we proceeded to checkerboard test the most synergistic and antagonistic clinical strains identified in our screen (Fig. 3 right). Strains DA81057 and DA81040 were most synergistic and antagonistic, respectively, for the combination of AMB-FLC; DA81050 and DA81109 were most synergistic and antagonistic, respectively, for the combination of ANI-FLC; and DA81027 and DA81024 were most synergistic and antagonistic, respectively, for the combination of AMB-ANI. FICi values for the clinical isolates with both methods were plotted against each other and fitted to a linear regression model (Fig. 3 right). The slope of the regression ranged from 1.066 to 1.317 (95% confidence interval, p = 0.001), indicating a strong agreement between the methods, though the combination plate tended to slightly overestimate FICi values compared to the checkerboard assay. Additionally, Bland-Altman analysis showed a low intrinsic bias (0.2) between the two methods, with 95% limits of agreement ranging from -0.67 to 1.09, further confirming the high level of concordance between results, despite the slight tendency of the combination plate to yield higher FICi values.

## Discussion

In this study, we present a new method for studying antifungal interactions. The combination plate allows testing of three antifungal combinations simultaneously and has been developed to provide overnight results. By replacing three checkerboard assays with a single, less labour-intensive, agar-based test, the combination plate provides an easier and faster approach to conducting antifungal interaction screenings, without compromising accuracy. For example, hands-on preparation time for one combination plate is approximately 5-10 minutes whilst one checkerboard may require upwards of60. Additionally, the method can easily be adjusted for the screening of isolates with varying susceptibility levels by simply altering the antifungal concentration in the loading chambers as needed.

Applying the combination plate to determine interactions in susceptible, tolerant or resistant fungal species could be clinically useful. This approach could help identify synergistic combinations against invasive infections, and development of new resistances could potentially be delayed. Perchance most worryingly is the increase in resistance towards echinocandins, found in both C. *glabrata* and C. *auris*. Combining empirical echinocandin treatment with other synergistic antifungals could prove to be an effective measure to widespread resistance emerging.

One of the primary goals with using this method was to obtain interaction results within an overnight incubation. Two key steps contributed to achieving this goal. First, we used YPD medium and agar throughout the process instead of RPMI. Even though RPMI medium is most commonly used in regular antifungal susceptibility testing, it has been argued that nutrient? deprivation from RPMI medium may lead to artificially low MIC results, as inhibited growth can be misinterpreted as susceptibility (40). For this assay, RPMI failed to support sufficient fungal growth for clear interaction patterns after overnight incubation, so YPD medium was used instead. While further standardization and optimization of media formulations for antifungal interaction testing is needed, results from the combination plates on YPD medium were similar to those obtained with RPMI in checkerboard assays, indicating that media choice has low impact on interaction results for this technology. The second step to reaching assay readability quickly was the use of an inoculated agar overlay. Using a suspension of cells in agar instead of streaking a liquid culture onto an agar plate enhanced fungal growth and produced a smooth lawn with distinct inhibition zones, aiding in the analysis of the combination plates.

In vitro studies of antifungal interactions have increased in recent years, as combination therapy is increasingly used to treat resistant pathogens. Traditionally, these studies have relied on checkerboard assays and typically involved smaller isolate collections (20,41–46). Our study was performed on a substantially larger isolate collection, and quantified 3 distinct combinations at once and on the same clinical collection. We observed striking differences between the interaction pattern of echinocandins compared to that of amphotericin and fluconazole.

Specifically, combinations involving an echinocandin displayed a variety of interaction profiles in our study, with close to 20% of strains exhibiting a positive interaction and 80% a negative one. This pattern, studies comprised of fewer isolates may not be able to capture. Indeed, varying responses to the same combinations are evident between studies (7,17), which has been posed to arise both due to the non-standardization of methodology and limited isolate collections (7). The isolate-specific variability in interaction pattern we observed here, makes a compelling argument for case-to-case testing of antifungal interaction in hospital clinics. Such routine clinical use of interaction testing of antifungals requires an easy to use and affordable test, which we think this combination plate can provide.

The combination of amphotericin B and fluconazole studied here showed, in contrast to the echinocandin combinations, limited variability and was overwhelmingly antagonistic, with all but one strain exhibiting antagonism. This combination is known to show discrepancies between in vivo an in vitro studies. Thus, the combination plate and other in vitro methodologies consistently report this combination as antagonistic (44,45,47), despite the favourable effects seen in vivo (48). A recent study showed that the difference seen may be due to concentration dependant interactions based on the free-drug levels in plasma (49). Deciphering these various physiological factors which may affect drug availability is necessary to help bridge the knowledge between in vitro and in vivo, aiding the tailoring of antifungal combination therapy regimens.

To categorize the interactions quantified in this study, we adhered to the original definition of the FIC index (33,34). Interactions with an index above 1 were classified as antagonistic, and those below 1 as synergistic. Some studies use alternative thresholds, such as an upper limit of 0.5 for synergy and a lower limit of 4 for antagonism, to reduce misclassification from the intrinsic variability in checkerboard assays (see below), though these thresholds lack clinical validation. Therefore, we chose to present our findings using a straightforward classification. We hope that large-scale studies enabled by the here proposed and similar technologies will ultimately help define clinically relevant thresholds for synergy and antagonism, though this goal remains a future step in method development.

Checkerboard assays, while common, have inherent difficulties in readability. They rely on serial dilutions, which have a margin of error of one well, leading to potential variability of up to four-fold in the FICi readout due to this inherent ambiguity. In contrast, the combination plate employs a continuous gradient of antifungals rather than serial dilutions, reducing variability as shown in the consistent FICi values from our screening.

Advances in the field of antifungal interaction testing have been made with the invention of several predictive chemogenomic methods(21–23). These methods hold great strength and can be valuable when exploring new combinations, developing new antifungals or evaluating interactions with other drugs. However, they may miss out on isolate-specific interactions which need to be tested on a case-to-case basis. Here the combination plate can serve as a complement to the in-silico methods to quickly evaluate if predicted synergies are present in clinical isolates, and if interaction profiles vary.

Although the combination plate is less labour-intensive and faster to perform, it maintains accuracy. Our cross-validation of the most synergistic and antagonistic drug-pathogen combinations with checkerboard assays demonstrates that results from this assay are directly comparable to those obtained from checkerboards, which are widely used but significantly more time-consuming to perform and interpret.

One limitation of the combination plate is that it primarily samples the concentration space near the MICs of the individual antifungal agents. Interactions between compounds, however, may vary at different concentrations (50,51), which a checkerboard assay could detect. We argue that the reduced inherent variability of the combination plate compensates for this limitation, especially as clinically relevant interactions are likely to occur around inhibitory concentrations, which is precisely where the combination plate operates.

In conclusion, the combination plate method presented here offers reproducible, overnight results when testing antifungal interactions in vitro. It is easily scalable, allowing larger screens as the one presented here which showed isolate-specific interaction profiles to the combinations tested. The current results highlight the need for individual testing prior to implementation of combination treatment, which could both improve the therapeutic outcome and slow down resistance emergence. Due to its simplicity the combination plate could readily be implemented in both clinical and research labs, to further advance the field of antifungal interactions.

## Materials and Methods

### Fungal isolates and growth conditions

For technical validation and method calibration the *Candida albicans* American Type Culture Collection (ATCC) strain 90028, obtained from the Department of Clinical Microbiology at Karolinska Institutet, was used. The fungus was grown overnight at 30°C, in liquid cultures using Yeast Peptone Dextrose (YPD) broth (Sigma-Aldrich, Ref. Y1375) in an orbital shaking incubator set at 190 rpm. All clinical isolates (n=92) were obtained from the Department of Clinical Microbiology of Karolinska Institutet, Huddinge. The isolates tested in this study were sampled from *C. albicans* bloodstream and vulvovaginal infections collected between 2021-2024 at Karolinska University Hospital, Stockholm, Sweden. Isolates were stored at - 80°C in freezing medium (-Nutrient broth (Oxoid CM0067) 21,25 g/L (2,13%)-Glycerol 85% (VWR 24384) 184,5 g/L (=150 mL/L) (12,75%)) until use and subsequently tested. Overnight cultures were prepared by transferring the yeast from frozen vials into 2 ml of YPD broth and incubated at 190 rpm orbital shaking.

### Antifungals

Antifungals were suspended in DMSO or water, per European Committee on Antimicrobial Susceptibility Testing (EUCAST) recommendation (52) to a final concentration of 10 mg/ml, aliquoted in 20 μl, and stored at -20°C until use. Antifungals used in this study include micafungin (Sigma-Aldrich, Ref. SML2268), anidulafungin (Sigma-Aldrich, Ref. SML2288), amphotericin B (ThermoFisher Scientific, Ref. J61491.03), fluconazole (Sigma-Aldrich, Ref. PHR1160) and voriconazole (ThermoFisher Scientific, Ref. 458670010).

### Broth microdilutions

Minimum inhibitory concentration (MIC) was determined for all antifungals for C. *albicans* ATCC strain 90028 using broth microdilution according to EUCAST guidelines (52) with minor modifications. RPMI medium with HEPES (Gibco, Ref. 130118031) was used instead of MOPS, plates were incubated at 30°C, and results were read by eye. Measurements were conducted with one biological replicate. Two-fold serial dilutions of the antifungals were prepared in 96-well round bottom microtiter plates, leading to a 100 μl suspension of the antifungals at 2X the final concentration. Wells were inoculated with 100 μl of yeast suspension prepared to 0.5 McFarland cells in RPMI medium. The MIC was defined as the concentration where 50% growth inhibition was seen (MIC50), except for amphotericin B where the 90% growth inhibition was deemed as the MIC (MIC90), in accordance to EUCAST guidelines.

### Optimization of fungal growth conditions

The six laboratory medium tested were:

1. BD BACTO^™^ Brain Heart Infusion (BHI) broth (Becton Dickinson, Ref. 237500)
2. BD DIFCO^™^ Mueller-Hinton (MH) broth (Becton Dickinson, Ref. 275730)
3. MH-II broth (Becton Dickinson, Ref. 212322)
4. Lysogeny Broth (LB) (Sigma-Aldrich, Ref. L3522)
5. Yeast Peptone Dextrose (YPD) broth (Sigma-Aldrich, Ref. Y1375)
6. RPMI medium (Gibco, Ref. 130118031)

2% D-glucose (w/v) was supplemented to all media, except YPD, and plates were prepared in-house with 1.5% agarose (w/v) (Sigma-Aldrich, Ref. A9639), except for YPD agar which was bought (Sigma-Aldrich, Ref. Y1500).

Three different plating methods were evaluated, namely:

1. Glass beads. 100 μl of undiluted fungal overnight culture was added directly to plates and spread using the beads.
2. Cotton swabbing, performed according to EUCAST guidelines but using undiluted overnight culture (53).
3. Inoculating an overlay using low-temperature gelling agarose. Overlay was prepared by mixing equal volumes of 3% (w/v) low-temperature gelling agarose (Sigma-Aldrich, Ref. A4018) with undiluted overnight culture, yielding a 1:2 dilution of the overnight. The low temperature gelling agarose was homogenized in a water bath set at 55°C.

### Combination plate calibration

Prior to the screening of clinical isolates, the concentrations of the antifungals to be used on the assay were calibrated using the ATCC type strain. MIC values obtained from the BMDs lay the basis for the calibration which was done for every antifungal. Optimal assay concentration was defined as the concentration where clear inhibition zones with defined edges formed as this aids the annotation of the key points during data analysis. During calibration, the three antifungal-loading reservoirs in a plate contained the same antifungal, but at different concentrations. Concentrations up to 600 times the MIC of each antifungal was tested, using an increment of either 10 (between 10-100X the MIC) or 100 (between 100-600X the MIC).

### Protocol validation and ATCC strain

After optimal growth conditions and antifungal concentrations were established for the assay, the developed protocol (supplementary protocol 2) was evaluated using the ATCC type strain. The strain was tested for the pairwise combination of ANI, AMB, and FLC in technical triplicates.

The antifungal-loading chambers were loaded with YPD agar containing one of the three antifungals. After the agar solidified, the assays were kept at 4°C for a minimum of 45 min, after which an underlay of 25 ml YPD agar was added. The agar was left the rest and solidify at room temperature for at least 2h before plating. Plating was done by using an inoculated agar overlay which was prepared by mixing equal volumes of a 3% low-temperature gelling agarose with a dense fungal overnight culture diluted 1:50 in fresh YPD medium, resulting in a final 1:100 dilution of the overnight culture. Methylene blue was added to the 3% agarose solution at a concentration of 1 μg/mL for a final concentration of 0.5 μg/ml

### Screening of clinical isolates

All clinical isolates (n=92) were screened for the pairwise combination of ANI, AMB, and FLC. Isolates were tested in technical triplicates of biological duplicates. Isolates were grown overnight as described and subsequently screened using the developed protocol (Supplementary protocol 2).

### Checkerboard technique

Checkerboard assays were performed for the ATCC 90028 C . *albicans* reference strain and the most synergistic and antagonistic clinical strains identified in the screen to verify the results. The checkerboard protocol used here was modified from the protocol published by Vitale *et*.*al* (54) and a step-by-step protocol is available in Supplementary protocol 1. Checkerboards were performed using 96-well round bottom microtiter plates in biological triplicates. The antifungal stocks prepared previously were suspended in RPMI medium at 4X the highest concentration to be used. The suspensions was 2-fold serially diluted and 50 μl of each antifungal, at 4X the final concentration, was added to their corresponding well. 100 μl of fungal suspension prepared at 0.5 McFarland was added to all experimental wells. RPMI medium served as a sterility control. After an overnight static incubation at 30°C the wells were resuspended and the optical density (OD) was measured at 450 nm using a Thermo Fisher Scientific Multiscan FC Type 357. For analyses, the OD values of the sterility control were subtracted from all experimental wells and the growth inhibition was calculated as a percentage by dividing the experimental wells with the growth control. To obtain the MIC values for the antifungals when used in combinations, points of theoretical additivity were picked. Based on the EUCAST method of MIC determination of yeasts in BMDs, the theoretical additivity is at 95% growth inhibition for the combinations of AMB-ANI and AMB-FLC whilst for the combination of ANI-FLC it is at 75% growth inhibition.

### Statistical analysis

Statistical analyses were performed using Graphpad Prism version 9, significance levels are noted at p=0.05 probability unless otherwise stated.

## Author contributions

Conceptualization: NFK, DA, VÖ, TUA

Formal analysis: NFK, HL

Funding acquisition: DA, VÖ

Investigation: NFK, HL, RS

Methodology: NFK, HL

Writing – original draft: NFK, HL

Writing – review and editing: all authors

## Funding

Funding was provided by grants awarded to VÖ and DIA …..Funders had no input in the investigation, analysis or decision to publish.

## Supplementary information

### Supplementary protocol 1: Checkerboard protocol for antifungal combinations

Protocol has been adapted from E. Dannouli et al and EUCAST protocol for MIC determination of yeasts by broth microdilution. The RPMI medium used throughout the protocol is supplemented with 2% glucose

1. Grow the fungal isolates of interest on YPD plates overnight at 30°C.
2. Prepare antifungal stocks in RPMI medium at 4X the highest concentration to be used in the assay.
3. Perform a 2-fold serial dilution in RPMI medium of each antifungal stock prepared, yielding a 4X dilution of all antifungal concentrations to be used.
4. Drug A is added to rows B-H. Add 50 ul of the prepared dilutions to each well with the highest concentration in row B and lowest concentration in row H to a 96-well round bottom microtiter plate.
5. Drug B is added to columns 2-11. Add 50 ul of the prepared dilutions to each well with the highest concentration in column 2 and lowest concentration in column 11.
  a. Based on step 3-4, wells B-H in column 1 contains only drug A and wells 2-11 in row A only drug B. Add 50 ul of RPMI medium to these wells for a final volume of 100 ul.
6. Prepare the fungal inoculum by suspending colonies in RPMI medium until a turbidity of 0.5 McFarland.
7. Add 100 ul of the fungal inoculum to all wells in column 1-11.
8. Well A1 serves as the growth control. Add 100 ul of RPMI medium to the 100 ul of inoculum
9. Well A12 serves as the sterility control. Add 200 ul of RPMI medium.
10. Incubate plate at 30°C overnight.
  a. To reduce evaporation, plates can be placed in plastic bags prior to incubation.
11. Analyse plates by measuring the OD at 450 nm
  a. If needed, resuspend wells prior to the OD measurement
12. Ensure sterility by comparing the sterility control with the empty wells in column 12.

### Supplementary protocol 2: Protocol for antifungal combination testing using combination plates

#### Day 1

1. Grow chosen strains in 2 ml of Yeast-Peptone-Dextrose (YPD) medium at 30°C in an orbital shaker overnight (ON).
  a. Places tubes diagonally in rack.
2. Prepare YPD agar according to manufacturer’s instruction, keep at 55°C.
3. Prepare a 1 mg/mL of methylene blue in sterile dH_2_O. Shield from direct light and store at 4°C until use.

#### Day 2

1. Dilute antifungal stock to the desired input concentration in YPD agar. Keep antifungal-agar in a water bath set at 55°C until use.
2. Load 0.5 mL of the antifungal agar to the assigned chamber of the CombiANT assay
  a. Ensure entire chamber is filled and remove any air bubbles.
3. Let agar solidify at room temperature, then keep at 4°C for at least 45 minutes until further use.
4. To activate the assay, remove plates from 4°C and add a 25 mL underlay of YPD agar to each plate.
  a. Add agar centrally of the assay, ensure all chambers are submerged and pipette away any bubbles.
  b. Do not move the plates until agar has solidified as it may disrupt the diffusion pattern
5. Let the agar set for at least 2 hours at room temperature after the last plate is cast
6. Prior to plating of the strains, prepare n + 1 mL (n = number of plates) of low-gelling temperature agarose at 3% (w/v) in sterile dH_2_O. Melt and keep agar at 55°C in a water bath.
  a. Add 1 μL of the methylene blue solution per mL of low-gelling temperature agarose solution prepared.
7. Dilute the ON cultures of the chosen strains 1:50 in YPD medium.
8. Per plate, mix 1 mL of the diluted ON cultures with 1 mL of the low-gelling temperature agarose with methylene blue. Mix by pipetting and pour immediately onto the plate.
  a. Ensure entire plate is covered by the overlay.
9. Let overlay solidify at room temperature for 10 min, then incubate ON at 30°C.

#### Day 3

1. Take pictures of the plates. Orient plates so chamber A is furthest down for easier analysis.
2. Identify the points of interest using the CombiANT Imager software (Rx Dynamics AB).
3. Input the data into the analysis algorithm.

## Notes

### Competing Interest Statement

NFK and DIA hold stock in Rx Dynamics AB. The company had no input in data interpretation, or decision to publish

### Summary of Updates

Ammended missing conflict of interest statement

